# The Fate of DIET Consortium exposed to continuous and intermittent electrochemical stimulation

**DOI:** 10.1101/2022.03.09.483589

**Authors:** Mon Oo Yee, Lars D.M. Ottosen, Amelia-Elena Rotaru

## Abstract

Electromethanogenesis is the electrochemical conversion of carbon dioxide (CO_2_) to methane (CH_4_) with microorganisms as catalysts, and it is a promising approach for biogas upgrading. Many studies have shown increased methane production by electrochemical stimulation, allegedly due to the enhancement of direct interspecies electron transfer (DIET) between electrogenic bacteria and methanogens. However, these studies were done with mixed cultures where the interspecies interactions were undefined, so the impact of an electrochemical stimulus on these interactions remains unresolved. Here we report that an electrochemical stimulus diminished the survival and activity of a defined DIET consortium. We investigated the impact of electrochemical stimulation on a co-culture of *Geobacter metallireducens* and *Methanosarcina barkeri*, where the two partners interact syntrophically via DIET. The electrochemical stimulus was a cathode poised at - 700 mV (versus the standard hydrogen electrode, SHE). Electrochemical stimulation was provided either continuously or intermittently, and so was the food substrate. Compared to the untreated DIET co-culture, all treatment strategies relying on cathodic current additions resulted in acetate accumulation, lower methane production and lower cell numbers for both partners, indicating a detrimental impact of electrochemical stimulation to a DIET consortium. These findings suggest that future bioelectrochemical technologies must guarantee the survival of the syntrophic partners during current additions.

## Introduction

Bioelectrochemical systems for converting CO_2_ from biogas to methane (biogas upgrading) have gained popularity as an alternative to conventional upgrading by H_2_-addition. Typically, the product of anaerobic digestion - biogas, is a mixture of CH_4_ (35-65%) and CO_2_ (15-50%) with traces of other gases which cannot be used as fuel unless the CH_4_ concentration is higher than 95% (v/v) (Aryal et al., 2018; Da Costa Gomez, 2013). However, typically harsh chemical treatments are required to remove impurities and upgrade CO_2_ from biogas, and such treatments ultimately lead to environmentally harmful CO_2_ emissions (Bauer et al., 2013). Thus, biogas upgrading to methane using renewable electricity can be beneficial by sustaining “carbon neutral” fuel production (Van Eerten-Jansen et al., 2012) while serving as a form of energy storage (Geppert et al., 2016).

Several studies have suggested that electrochemical stimulation enhances interspecies interactions in anaerobic digesters, specifically direct-interspecies electron transfer (Lin et al., 2019; Zhao et al., 2015b, 2015a). Direct Interspecies Electron Transfer (DIET) is a mutualistic relationship where the two partners exchange electrons. During DIET, the syntroph oxidizes a reduced food substrate and donates electrons to a methanogen via a network of pili and multiheme cytochromes. The methanogen then uses electrons from the syntroph’s oxidative metabolism to convert carbon dioxide to methane (Rotaru et al., 2021). DIET interactions have been demonstrated between *Geobacter* sp. and *Methanosarcinales* methanogens (*Methanosarcina* sp., *Methanothrix* sp.) (Rotaru et al., 2014a, 2014b; Yee et al., 2019), which are abundant in some anaerobic digestors (De Vrieze et al., 2012; Shrestha et al., 2014; van Haandel et al., 2014).

Although some studies with mixed communities from anaerobic digesters suggested that electrochemical stimulation enhances DIET, another study reported contradictory observations noting that the addition of electricity inhibited DIET interactions and instead promoted hydrogenotrophic methanogenesis (Lee et al., 2017). In these studies, the interspecies relationships in the mixed community were speculative, and cathode potentials were undefined as the authors applied a voltage across the electrochemical cell. Thus, the impact of electrochemical stimulation on a DIET consortium has never been reported.

In bioelectrochemical systems, it has been shown that lower cathode potentials are more beneficial for increased methane yield. Only a few methanogens can retrieve electrons directly at a high cathodic potential (−400mV) when H_2_ does not form at the electrode abiotically (Beese-Vasbender et al., 2015; Yee et al., 2019; Yee and Rotaru, 2020). Conversely, at cathodic potentials below −500 mV vs. SHE, H_2_-consuming methanogens produce more methane when the cathodic potentials were lowest (Batlle-Vilanova et al., 2015; Cheng et al., 2009; Fu et al., 2015; Rosenbaum et al., 2011; Villano et al., 2010; Zhen et al., 2015b), likely due to methanogenesis from electrochemical H_2_ (Lohner 2014; Deutzmann 2015) or enzymatically-generated H_2_ (Deutzmann et al., 2015).

Since lower cathodic potentials are likely to be used in future applications for biogas upgrading, we investigated the effect of a low cathode potential (−700 mV vs. SHE) on a defined DIET co-culture of *G-metallireducens* and *M. barkeri*. To mimic the operational conditions of an anaerobic digestor, we tested representative CO_2_:CH_4_ headspace concentrations (50:50, v:v) and different feeding and current addition strategies. We tested various parameter changes, including continuous feed with substrate and electrical current, continuous substrate feed with intermittent current additions, and switching between the substrate and electrical current feed. Then, we monitored changes in substrate consumption, current draw, product formation and cell distribution. We showed that electric current addition negatively impacted methane production under all conditions tested. Additionally, we show that a high CO_2_ in headspace was favorable to hydrogenotrophic methanogenesis indicating that *Methanosarcina* decoupled from its DIET partner switching to a hydrogenotrophic metabolism. The decoupling of the two DIET partners was also supported by a significant drop in *Geobacter* cell numbers.

## Material and Methods

### Cultivation conditions

*Methanosarcina barkeri* MS (DSM 800) and *Geobacter metallireducens* GS-15 (DSM 7210) were originally purchased from the German culture collection DSMZ. We previously paired a co-culture of the two species in the presence of granular activated carbon (25 g/L) (Yee et al., 2019), and in this study we used the fifth transfer of this adapted co-culture. Growth was carried out in a previously reported basal medium (Yee et al., 2019). All cultivations were done under strict anaerobic and sterile conditions.

### Bioelectrochemical reactor setup and measurements

Two-chambered H cell reactors were setup as previously reported without stirring, to counteract noisy electrochemical signals (Yee et al., 2019). Briefly, the two chambers each with a total volume of 650 mL (Adams and Chittenden Scientific Glass, USA) were separated by a Nafion^™^ proton exchange membrane (Ion Power). Graphite rods with dimensions of 2.5 × 7.5 × 1.2 cm were used as both the working and counter electrodes and as reference electrodes we used leak-free Ag/AgCl (3.4 M KCl) (CMA Microdialysis, Sweden). The Ag/AgCl electrodes from CMA Microdialysis has a potential difference of +240 mV against the standard hydrogen electrode as reported by the manufacturer and confirmed in our laboratory. All potentials in this paper are presented against the standard hydrogen electrode (SHE) after conversion. To apply defined cathodic potentials and carry out chronoamperometry we used a multi-channel potentiostat (Multi Emstat, Palmsens, The Netherlands). The medium used was the modified DSMZ 120c medium (as mentioned above) and a reduced amount of GAC (2 g/L) was added into the working electrode chamber before autoclaving. The coculture at late-exponential growth (110 mL) was inoculated anaerobically to each reactor with a medium volume of 440 mL.

Three different treatment strategies were applied (**Fig. 1**), namely: (1) Feed and *continuous* current, (2) Feed and *intermittent* current, (3) Either feed or current. A control treatment was provided only with feed but without electricity addition (open circuit control). Ethanol was used as “feed” and maintained at concentrations of 1 to 2 mM, by additions whenever necessary. During “current” mode, the cathodes were poised at – 700 mV (vs. SHE). Intermittent current addition was carried out every 5-days.

**Figure 1.**
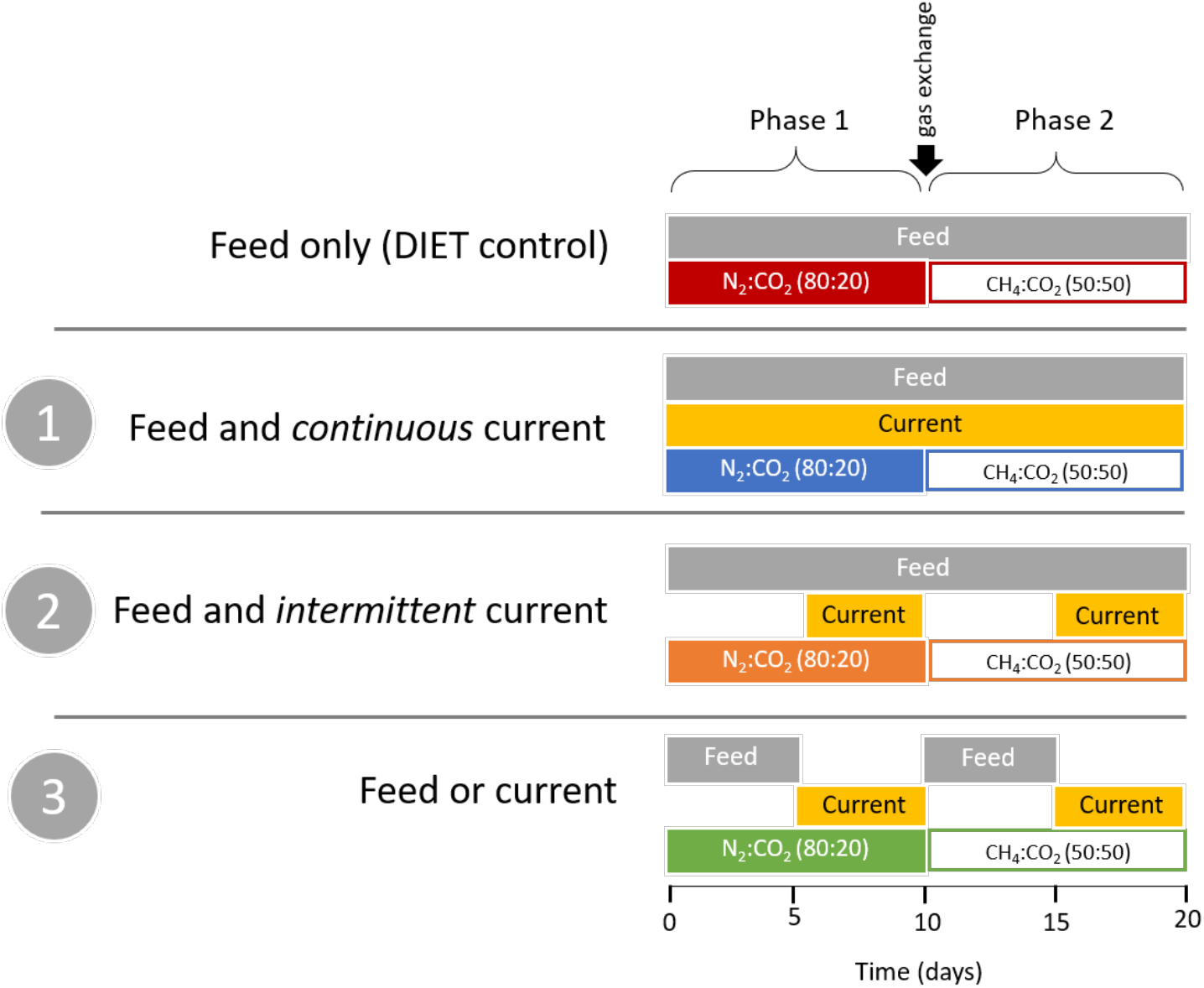
Diagram of treatment strategies applied to bioelectrochemical reactors testing the bioelectrochemical response of DIET co-cultures of *G. metallireducens/M. barkeri*. “Feed” corresponds to the maintenance of ethanol at 1 mM and “current” corresponds to a poised cathode at – 700 mV (vs. SHE).

During the first 10 days (phase 1) all incubations were done under an atmosphere with 80:20 N_2_:CO_2_ in the headspace. Afterwards (phase 2), the atmosphere was changed to a biogas mixture of CH_4_:CO_2_ (50:50) by a 20 minutes gas exchange and incubations were continued for another 10 days. Four replicate reactors were set up for each treatment condition but due to technical failures, only duplicate and triplicate reactors were carried out for the feed only treatment and for the feed and intermittent current treatment respectively. Abiotic reactors were also carried out for the different current treatments with ethanol added only at the start.

### Analytical measurements

Methane (CH_4_), hydrogen gas (H_2_) and ethanol concentrations were measured by gas chromatography, whereas acetate concentrations were measured by ion chromatography as we have previously described (Yee et al., 2019).

### DNA extraction

For DNA extraction, 20 ml sample were extracted anaerobically using hypodermic syringes and needles from the side ports of the bioelectrochemical reactors. To determine the cell numbers at the start of the incubations the co-culture inoculum was sampled (20 mL) with four technical replicates prior to inoculation into the reactors. From all 20mL samples, cells were pelleted at 5000 g for 15 min at 4C (Eppendorf), then pellets were frozen prior to DNA extraction. DNA extraction from pellets was done using MasterPure^™^ DNA purification kit (Epicentre, US) according to the manufacturers protocol. The resulting DNA was then eluted in 35 μL TE buffer.

### Preparation of standards for qPCR

The genomic extracts of *G. metallireducens* and *M. barkeri* were obtained similarly as described under the previous section. There were used as template DNAs for making the standards. For detecting *G. metallireducens*, primers specific for *Geobacteriaceae* 16s rRNA (GEO_494F; 5’ AGGAAGCACCGGCTAACTCC 3’) and (GEO_ 825R; 5’ TACCCGCRACACCTAGT 3’) were used (Holmes et al., 2002). For *M. barkeri*, the *Methanosarcinaceae* 16s rRNA specific primer pairs (MSC_380F; 5’ GAAACCGYGATAAGGGGA 3’) and (MSC_828R; 5’ TAGCGARCATCGTTTACG 3’) were used (Yu et al., 2005). PCR reaction was prepared in 50 μl final volume with 30.75 μl PCR-grade water, 5 μl of dNTP (2 mM), 5 μl of MgCl_2_ (25 mM), forward and reverser primer at 1 μl each, 5 μl Taq polymerase buffer (10x), 1 μl of genomic DNA as template, 1 μl of bovine serum albumin (BSA) and 0.25 μl of Taq polymerase (5U/ μl) (Invitrogen). The PCR amplification was as follow: 1) hot-start at 94°C for 10 mins, 2) denaturation at 94°C for 45 sec, 3) annealing for 45 sec at 55°C for *Geobacter* primers and at 50°C for *Methanosarcina*, 4) extension at 72°C for 45 sec, after which steps 2 to 4 were repeated for 35 cycles. The final extension step was at 72°C for 10 mins.

The PCR amplicons were purified in 1.2% agarose in TAE buffer and gel extracted with QIAEX II gel extraction kit following manufacturer’s protocol. The purified PCR product was ligated into pCR^™^4-TOPO^™^ vector from TOPO^™^ TA Cloning^™^ Kit and transformed into Top10 chemically competent *Escherichia coli* cells (Thermo-Fisher Scientific) following the manufacturer’s instructions. Transformed cells were grown and selected on LB plates with both ampicillin (100 μg/ml) and kanamycin (50 μg/ml). Colonies were picked and screened by PCR amplifications under the same conditions as described above. Plasmids were extracted from successful colonies with Miniprep Qiagen plasmid extraction kit. PCR was repeated with the plasmid as the template and the product was sequenced for verification. Successful plasmids were quantified with Quant-iT^™^ PicoGreen^™^ dsDNA Assay Kit (Invitrogen) and diluted from 10^8^ copies to 100 copies. These were used as standards for qPCR.

### Conditions for qPCR

The same primer pairs were used to prepare qPCR reactions. qPCR mix was prepared in a final volume of 25 μl of which contained 12.5 μl of RealQ Plus 2x master mix (Denmark), 0.5 μl of specific forward or reverse primer (20 μM) and 1 μl template. The PCR amplification reaction was as follows: 1) initial denaturation at 95°C for 15 mins, 2) denaturation at 95°C for 30 sec, 3) annealing for 30 sec at 55°C for *Geobacter* primers or 50°C for Methanosarcina primers, 4) extension at 72°C for 30 sec. Step 2 to 4 were repeated 40 times. Before termination, the PCR sample was treated at 95°C for 5 mins to obtain the melt curve.

## Results and discussion

DIET co-cultures of *G. metallireducens* and *M. barkeri* were exposed to six different conditions (**Fig. 1**), including different current addition strategies (constant or intermittent), feed strategies (constant or intermittent), and a different gas-atmosphere to identify the best strategy for electrochemical stimulation. The first 10 day-period (start-up period) was under an atmosphere typical of methanogenic incubations in the laboratory (80:20 N_2_: CO_2_). In the second 10-day period, the headspace-atmosphere was replaced with artificial biogas (50:50 CH_4_: CO_2_ gas mixture). To mimic organic load (feed) from anaerobic digestion, we maintained ethanol at 1 mM by supplementing every time there was a drop below the 1 mM mark, except for the one treatment (#3, feed or current) in which ethanol was added only at the beginning of the test.

We first tested the viability of a DIET co-culture (*G. metallireducens – M. barkeri*) in bioelectrochemical reactors under open-circuit conditions without a poised cathode. These co-cultures were fed continuously with a DIET substrate, ethanol (>1 mM) and provided with a reduced amount of granular activated carbon, which is known to stimulate their growth. These DIET consortia did not produce detectable H_2_ in the headspace of the reactors (**Fig. 2a**), consistent with a H_2_-independent interaction between *Geobacter* and *Methanosarcina*. When DIET consortia were exposed to a biogas atmosphere of 50:50 CH_4_:CO_2_ instead of the typical 80:20 N_2_:CO_2_ atmosphere, we observed a 3-fold increase in methanogenesis rate (**Fig. 2b**). Increasing methanogenesis rates did not match the ethanol oxidation rates, which decreased to half under the biogas atmosphere (p=0.005, **Fig. 2d**). Also, acetoclastic methanogenesis or perhaps syntrophic acetate oxidation appeared to be favored under this atmosphere because there was an apparent net consumption of 1.35 mM acetate, whereas 2 mM acetate accumulated under a typical N_2_:CO_2_ atmosphere (**Fig. 2e**). From previous observations (Rotaru et al., 2014b; Yee et al., 2019) and according to stoichiometry, we expected that each mol ethanol would transiently generate one mol acetate during the initial stages of a DIET-co-culture. Indeed, under a typical N_2_:CO_2_ atmosphere, the cocultures showed almost a 1:1 ratio of acetate to ethanol that dropped circa 3-fold under a biogas atmosphere. Both species in the DIET co-cultures increased in numbers in these open circuit reactors (**Fig. 2c**). *G. metallireducens* increased circa 8-fold and maintained cell numbers independent of the change in the atmospheric gases of the reactors, indicating a biogas atmosphere was not detrimental to their cell growth. However, it was detrimental to *Geobacter*’s ability to carry out ethanol oxidation which was halved (**Fig. 2d**). *M. barkeri* cell numbers increased 12-fold under a N_2_:CO_2_ atmosphere and were expected to increase under a biogas-atmosphere to match the 3-fold higher methanogenesis rates. However, the cell numbers increased insignificantly compared to the previous phase (1.5-fold increase; p=0.12). The ratios of *Geobacter:Methanosarcina* in a typical DIET-co-culture are 1:6 (Rotaru 2014a), and in this open circuit bioelectrochemical system, the ratio was close 14:1 (phase 1) and 10:1 (phase 2) of *Geobacter* to *Methanosarcina*.

**Figure 2.**
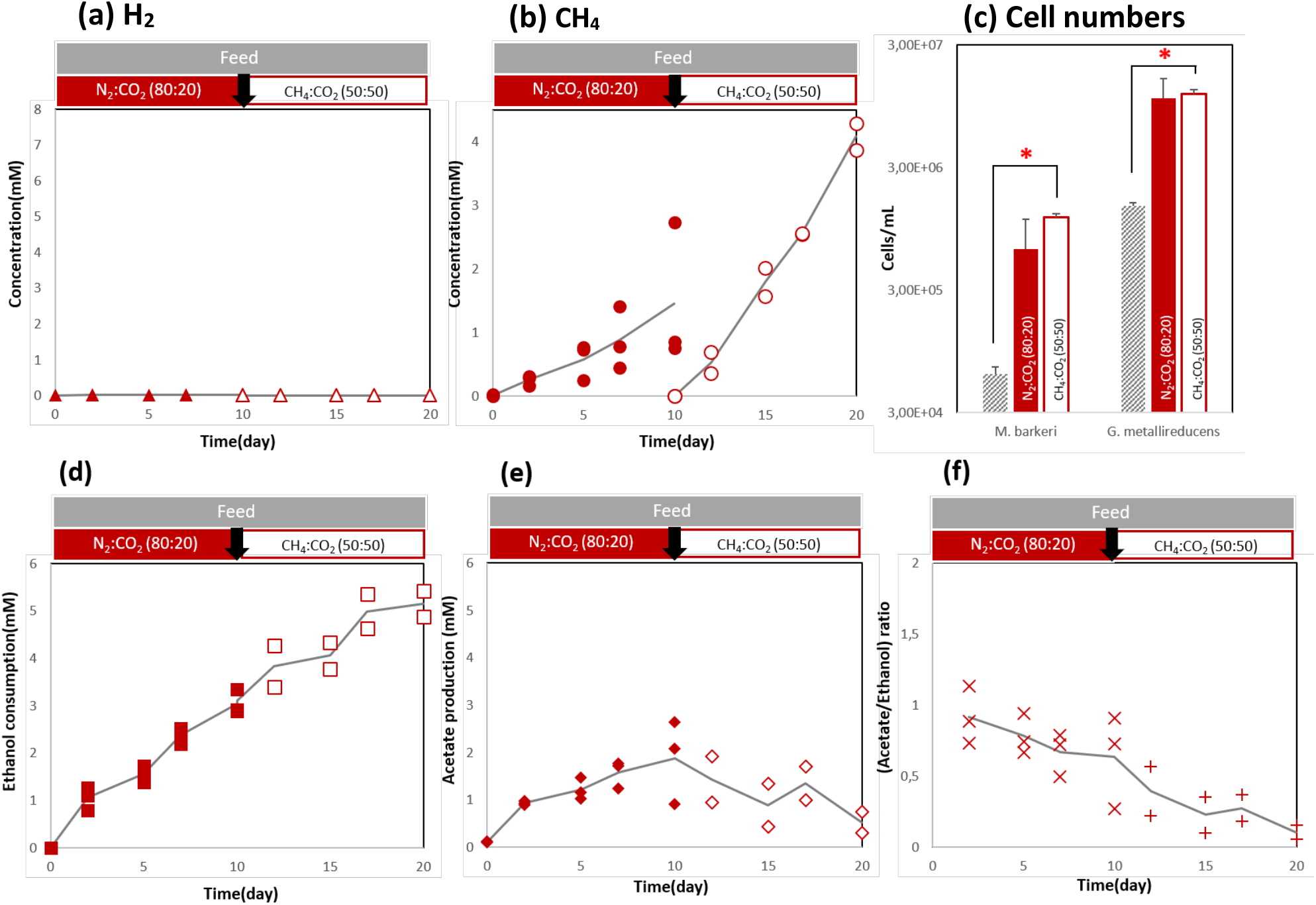
DIET co-culture of *G. metallireducens* and *M. barkeri* established in open circuit bioelectrochemical reactors with disconnected cathodes. (a) H_2_ production, (b) methane production above background, (c) cell numbers obtained by qPCR, (d) cumulative ethanol consumed, (e) acetate production, (f) ratio of acetate to ethanol. (n=3) in the first cycle, (n=3) in the second cycle. The treatment strategy is depicted above the graph. The black arrow depicts time of gas exchange. The grey line is the average of replicates, closed symbols and crosses are measured under a headspace of N_2_: CO_2_ (80:20), the open symbols and pluses are measured after atmosphere exchange to biogas (CH_4_: CO_2_ 50:50).

### Continuous feed and electrochemical stimulation led to lower methanogenic rates and decoupled DIET cocultures

One strategy to store electrical energy into a fuel like methane would be to supply a poised cathode directly to an operational anaerobic digestor while supplying a standard organic feed to maintain the growth of syntrophic consortia (**Fig. 1**). However, electrical current additions may effectively inhibit the syntrophic bacteria since the methanogen would have a competitive alternative electron donor – the poised cathode instead of the syntrophic partner. Likewise, it has been previously shown that H_2_-producing syntrophic partners have been effectively inhibited by adding high H_2_ concentrations, a competitive electron donor for hydrogenotrophic methanogenesis (Stams et al., 2012). Here we tested whether *Geobacter* would be negatively impacted during continuous electrochemical stimulation with a cathode at −700mV as an alternative electron donor for *M. barkeri*.

During the first phase (N_2_:CO_2_), the continuous cathodic electron supply at – 700 mV (vs. SHE) led to the evolution of ca. 0.6 mM/day of H_2_ (first phase, **Fig. 3a**), in stark contrast to undetectable H_2_-evolution in the continuous ethanol fed DIET co-cultures (**Fig. 2a**). The addition of a poised cathode that leads to abiotic H_2_ accumulation (**Fig. 1S**) did not immediately favor hydrogenotrophic methanogenesis by *M. barkeri*, likely because *Methanosarcina* is not adapted to H_2_-utilization when growing via DIET. The metabolically versatile *M. barkeri* exhibits diauxic growth in pure culture and uses preferentially H_2_ over acetate (Hutten et al., 1980; Smith and Mah, 1978). It was therefore surprising that a *M. barkeri* adapted to DIET electron uptake form a *Geobacter* was unable to switch to hydrogenotrophic methanogenesis quickly. Under a biogas atmosphere (50:50 v:v CH_4_:CO_2_), H_2_-evolution rates decreased by 6-fold compared to under a typical co-culture atmosphere of N_2_:CO_2_ (**Fig. 3a**). At the same time, methanogenesis rates increased, predictably, due to an increase in hydrogenotrophic methanogenesis (**Fig. 3b**). In contrast, H_2_ evolution rates doubled under the biogas atmosphere in cell-free abiotic reactors (**Fig. 1S**). Thus, we ruled out the possibility that the decrease in H_2_ was due to an electrochemical limitation under a biogas atmosphere.

**Figure 3.**
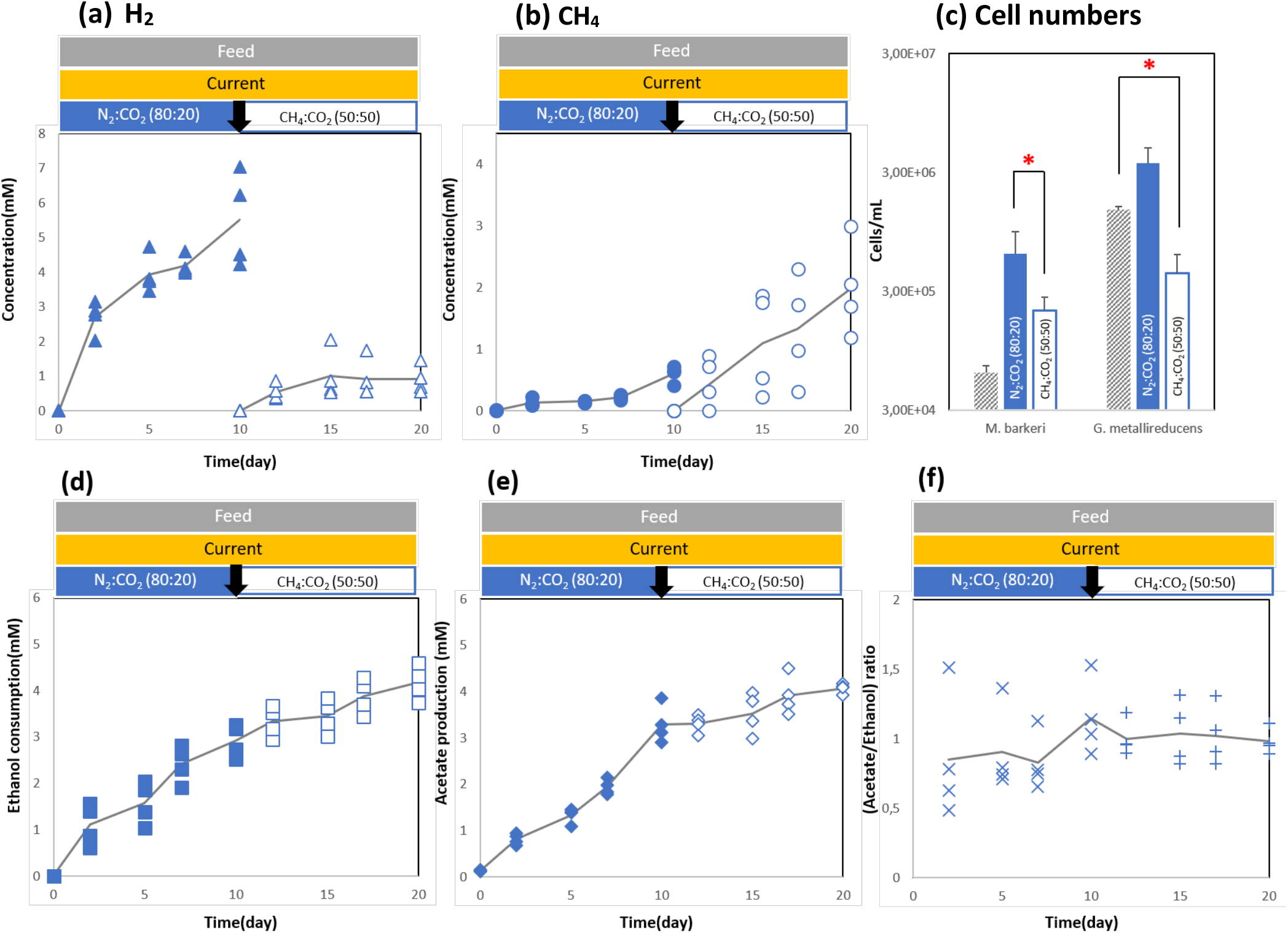
A DIET co-culture of *G. metallireducens* and *M. barkeri* exposed to a continuous bioelectrochemical stimulus - a poised cathode at – 700 mV (vs. SHE) (n=4) (a) H_2_ production, (b) cumulative methane production above background (c) cell numbers obtained by qPCR; (d) accumulated ethanol consumption; (e) acetate production; (f) molar ratio of acetate to ethanol. The treatment strategy is depicted above the graph. The black arrow depicts time of gas exchange. The grey line is the average of replicates, closed symbols and crosses are measured under a headspace of N_2_: CO_2_ (80:20), the open symbols and pluses are measured after flushing with CH_4_: CO_2_ (50:50). A representative current accumulation is described in Fig. 2S.

In contrast to DIET controls, the continuous feed and cathodic electron supply did not promote methanogenesis by continuously stimulated co-cultures neither under a N_2_:CO_2_ atmosphere (2-fold drop; p=0.14) **Fig. 3b**) nor under a biogas-atmosphere (2-fold drop; p=0.004).

Surprisingly ethanol oxidation was not impacted by the addition of a poised cathode at −700mV (**Fig. 3d**) since ethanol oxidation proceeded at identical rates (0.3 ± 0.03 mM/day) to those observed in DIET controls (**Fig 2d**) under a typical co-culture atmosphere. But ethanol oxidation rates dropped significantly (0.1 ± 0.007 mM/day) under a biogas atmosphere. This is because the methanogen got decoupled from its partner switching to hydrogenotrophic methanogenesis (observed H_2_ decrease and methane increase) instead of DIET. This decoupling was also confirmed by acetate accumulating twice as fast in continuously stimulated co-cultures (**Fig. 3e**; 0.32 ± 0.04 mM /day; n=4; p<0.05) in contrast to DIET-controls (0.17 ± 0.09 mM/day; n=2) (**Fig. 3e**). Acetate accumulation continued at quarter speed of the previous phase (Fig. 3; 0.08 ± 0.04 mM/day; n=4; p=0.04) under a biogas atmosphere in contrast to being consumed during DIET (Fig. 2; - 0.2 ± 0.07 mM/day; n=2). Independent of the gas-atmosphere when a continuous electrochemical stimulus was applied, the ratio of acetate to ethanol remained 1:1, clearly showing that acetate could not be metabolized in the presence of a cathode at −700 mV (**Fig. 3f**).

During incubation with an N_2_:CO_2_ atmosphere and continuous stimulation (cathodic and feed), *Geobacter* cell numbers increased by 2.5-fold and *Methanosarcina* by 10-fold compared a DIET-co-culture incubated with GAC, reaching a typical DIET co-culture distribution of 1:6 of *Methanosarcina* to *Geobacter* (Rotaru et al., 2014a). Both cell types decreased when switching to a biogas atmosphere, with *M. barkeri* showing a 3-fold decrease in cell numbers (**Fig. 3c**), and *Geobacter* 9-fold (**Fig. 3c**). Nevertheless, under a biogas atmosphere, H_2_-did not accumulate, whereas acetate did accumulate, suggestive of a switch in *M. barkeri’s* metabolism towards hydrogenotrophic instead of DIET and acetoclastic methanogenesis. A higher CO_2_ partial pressure may have decoupled the DIET partnership, but other parameters may be involved (e.g., pH, CH_4_, N_2_ etc.) as well, which remain to be tested.

### Intermittent additions had better outcomes for the survival of DIET consortia

The intermittent addition of electrical current is a feasible strategy for biogas upgrading because green energy resources (solar, wind, waves) are sporadic demanding strategies for energy storage. Here we tested how intermittent electrical current additions impact a DIET-co-culture (Methanosarcina-Geobacter) fed continuously with its substrate - ethanol. As noted above, continuous current and feed addition decoupled the DIET association, providing a strong selective pressure favorable for the methanogens and suppressing *Geobacter*.

Under intermittent current additions (−700mV on/off every 5 days) the co-culture was performing worse than under DIET conditions both in methane production and ethanol utilization. However, the *Methanosarcina* cell numbers remained unaffected by intermittent current additions (p=0.09), whereas *Geobacter* cell numbers still decreased (phase 1: 2-fold, p=0.04; phase 2: 13-fold, p<0.00001).

Unlike continuous current additions, under intermittent current additions we observed:

- 2 to 3-fold lower methane production (**Fig. 5b**, p<0.05)
- similar or a slightly decreasing ethanol consumption (1.18-fold) (**Fig. 5d)**
- similar acetate and H_2_ production profiles indicative of non-hydrogenotrophic methanogenesis during the N_2_:CO_2_-phase; however due to lower current input during intermittent additions, H_2_ production rates were 3-fold lower (p=0.04) (**Fig. 5a & e**)
- similar acetate and H_2_ production profiles during the CO_2_:CH_4_-phase indicative of adaptation to hydrogenotrophic methanogenesis; however, acetate accumulation rates were 3.5-fold lower than in co-cultures continuously exposed to −700mV. Lower acetate accumulation is likely due to less H_2_-accumulating during intermittent current additions. H_2_ is known to be consumed before acetate-utilization starts in *Methanosarcina barkeri* (Ferguson et al., 1983), therefore being exposed to less H_2_ may be more favorable to be less detrimental to acetoclastic methanogenesis.
- *Methanosarcina* cell numbers were highest during intermittent current additions, in contrast to continuous current additions when they were 3-fold lower. Although *G. metallireducens* showed a drop in cell numbers during intermittent additions (phase 1: 2-fold, p=0.04; phase 2: 13-fold, p<0.00001), it was to a much lesser extent than it did during continuous current additions (phase 1: 3-fold, p=0.03; phase 2: 28-fold, p<0.00001). Collectively, these results suggest less detrimental impact when electricity is supplied intermittently, and more detrimental when supplied continuously.

**Figure 5.**
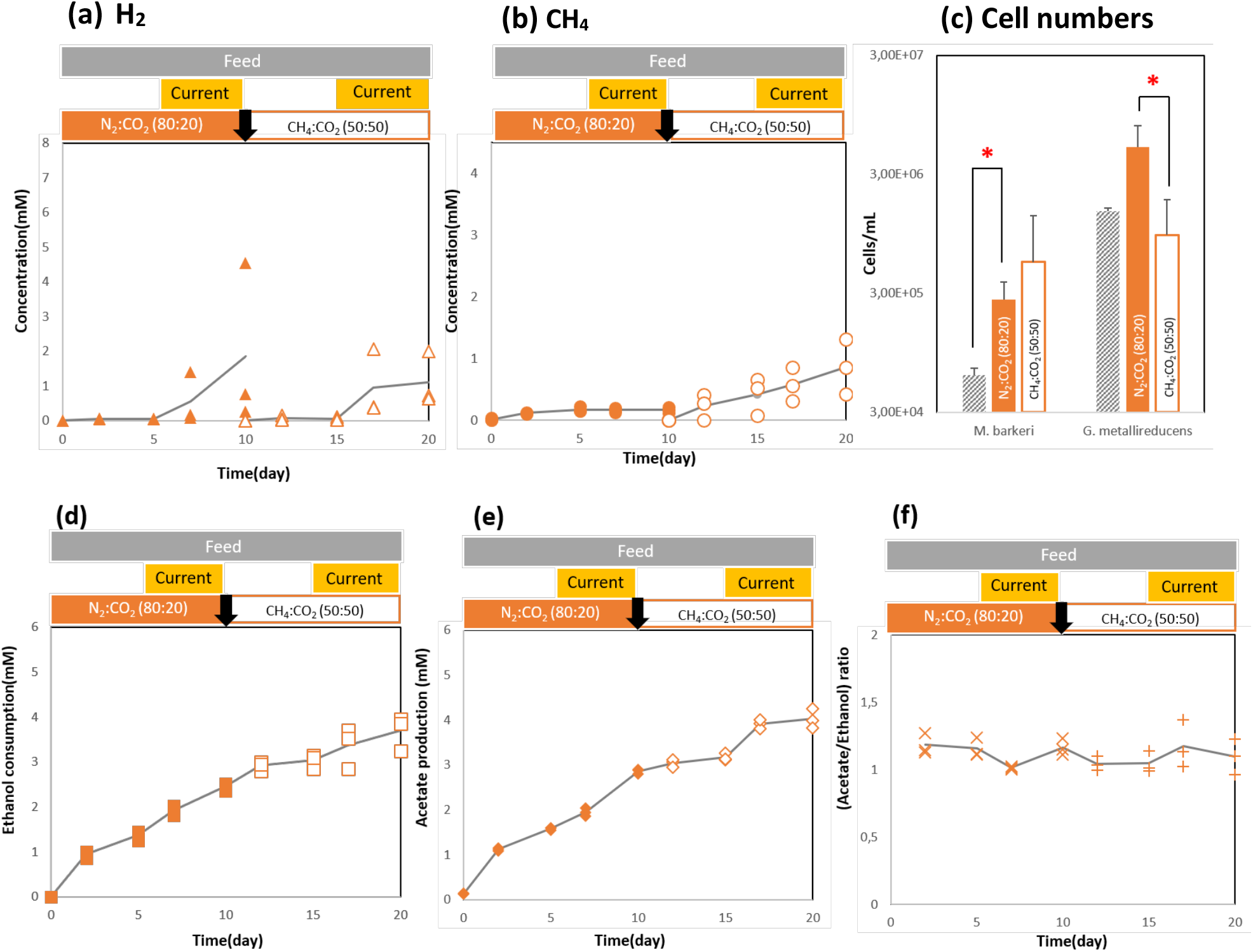
Co-culture of *G. metallireducens* and *M. barkeri* established in bioelectrochemical reactors with a cathode poised intermittently at – 700 mV (vs. SHE) (n=3) (a) H_2_ production, (b) methane production above background (c) cell numbers obtained by qPCR; (d) accumulated ethanol consumption; (e) acetate production; (f) molar ratio of acetate to ethanol. The treatment strategy is depicted above the graph. The black arrow depicts time of gas exchange. The grey line is the average of replicates, closed symbols and crosses are measured under a headspace of N_2_: CO_2_ (80:20), the open symbols and pluses are measured after flushing with CH_4_: CO_2_ (50:50). A representative current accumulation is described in Fig. 2S.

For the last treatment strategy (**Fig. 6**), we alternated feed with electric current additions at 5-day intervals. The objective for this treatment was to investigate if *Geobacter* could survive starvation periods (electron donor limitation) when the sole electron donor available - a cathode - benefits only the methanogen (Gregory et al., 2004; Lovley et al., 2011). Under a N_2_:CO_2_ atmosphere, *Geobacter* cell numbers decreased 2.4-fold compared to DIET-control co-cultures, but were comparable to those observed in the other two treatments with electrochemical stimulation. On the other hand, under the biogas phase, when the methanogen adjusted to utilizing the electrode at −700mV as preferable electron donor, *Geobacter-cell* numbers dropped 37-fold compared to those in DIET-co-cultures. This decrease in cell numbers is the lowest observed when looking at all other treatments with electrochemical stimulation (28-fold for continuous and 13-fold for intermittent). We thus show that *Geobacter* was negatively impacted by substrate limitation and competitive processes providing electrons for the methanogen. Nevertheless, even in low numbers, the remaining *Geobacter* recuperated quickly once it was provided with food again, indicating their population can persist substrate limitation and detrimental cathodic current additions.

**Figure 6.**
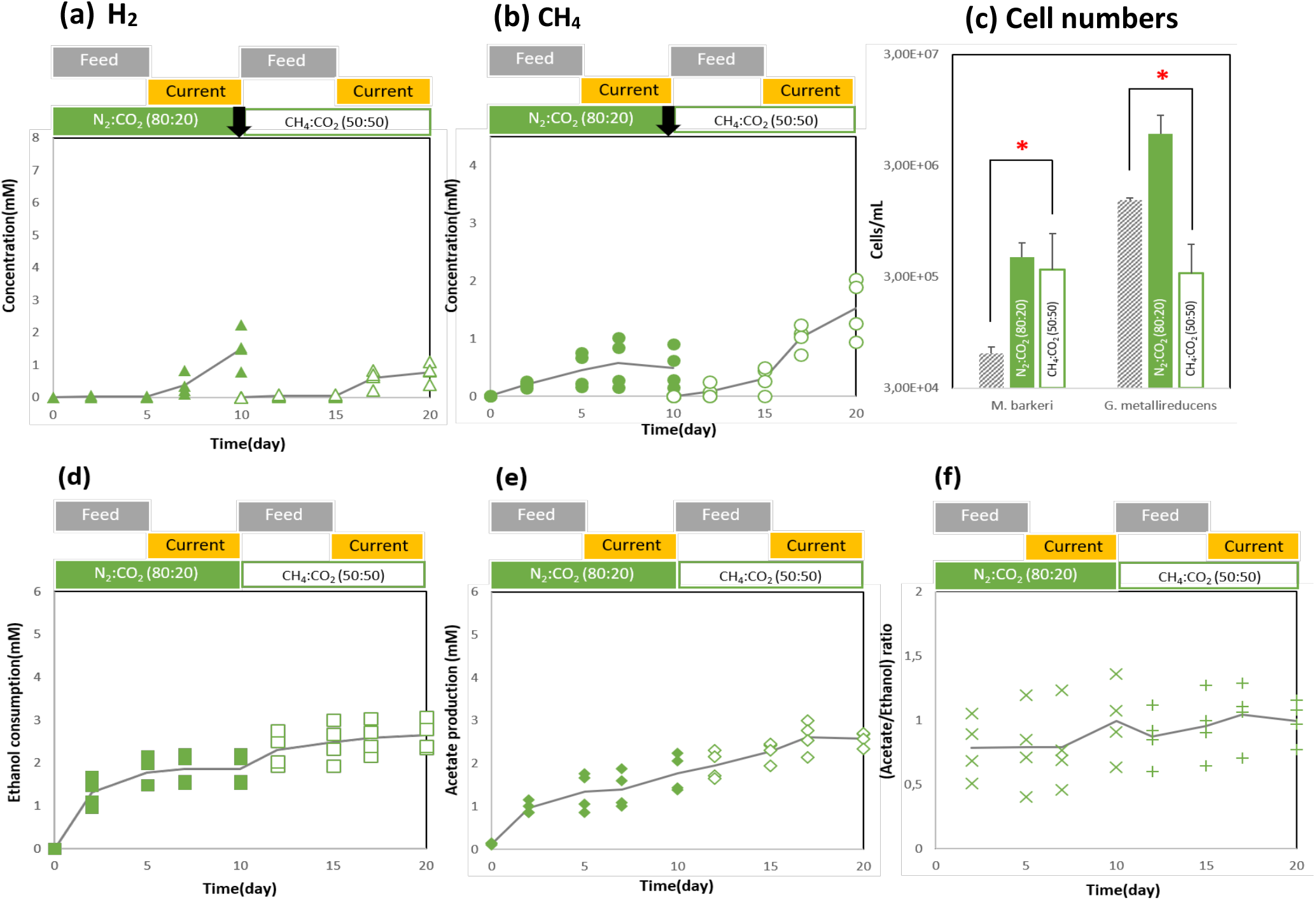
Co-culture of *G. metallireducens* and *M. barkeri* established in bioelectrochemical reactors fed with ethanol or poised intermittently with a cathode at – 700 mV (vs. SHE) (n=3) (a) H_2_ production, (b) methane production above background (c) cell numbers obtained by qPCR; (d) accumulated ethanol consumption; (e) acetate production; (f) molar ratio of acetate to ethanol. The treatment strategy is depicted above the graph. The black arrow depicts time of gas exchange. The grey line is the average of replicates, closed symbols and crosses are measured under a headspace of N_2_: CO_2_ (80:20), the open symbols and pluses are measured after flushing with CH_4_: CO_2_ (50:50). A representative current accumulation is described in Fig. 2S.

## Implications

This study investigated the effects of electricity addition to DIET interactions in an artificial consortium (*G. metallireducens* and *M. barkeri*) and subjected it to different feeding and electricity addition strategies. Considering the overall methane production and cell numbers at the end of the experiment, we found that the DIET only control, with no current additions, had the best output. The addition of a cathode poised at – 700 mV resulted in H_2_ production but was only consumed by *M. barkeri* under a 50% headspace CO_2_ concentration. In anaerobic digesters producing biogas, headspace concentrations of CO_2_ are often high (up to 50%) (Aryal et al., 2018). This meant that the introduction of H_2_ via direct injections or via an electrode could potentially cause the DIET-methanogens like *M. barkeri* to switch to H_2_ utilization. However, dominant methanogens in anaerobic digesters belong to the genus *Methanothrix* (Rotaru et al., 2014b), which are unable to utilize H_2_, and thus their DIET interactions are unlikely to be impacted in such a case. This was supported by studies that documented an enrichment of methanogenic DIET consortia of *Geobacter* and *Methanothrix* when anaerobic digestor inocula were supplied with a poised electrode (Lin et al., 2019; Zhao et al., 2015b, 2015a). However, these studies did not measure the cathode potential or H_2_ build-up.

Headspace CO_2_ limitation had been studied in the context of biogas upgrading in mixed hydrogenotrophic methanogenic cultures and a reduction and total inhibition of H_2_ consumption was found to be at CO_2_ concentrations below 12% and 6%, respectively (Agneessens et al., 2017; Garcia-Robledo et al., 2016). For a pure hydrogenotrophic methanogen, *Methanobacterium congolense*, the CO_2_ threshold for methanogenesis during biogas upgrading can be as low as 2% (Chen et al., 2019). The H_2_ threshold of strictly hydrogenotrophic methanogens is also magnitudes lower than generalists such as *M. barkeri* (Kral et al., 1998; Lovley, 1985). Thus, these differences in CO_2_ and H_2_ thresholds of methanogens might also explain why mostly hydrogenotrophic were often enriched in bioelectrochemical systems as studies were often carried out with CO_2_ concentrations around 20-30% and most likely with incomplete diffusion limited mass transfer to the relevant microbial communities (Cheng et al., 2009; Siegert et al., 2014, 2015; Van Eerten-Jansen et al., 2013). Similarly, headspace composition had also been shown to influence product profiles in a sulphate-reducing electrochemical reactor (Sharma et al., 2014). We suggest that further tests for biogas upgrading coupled to anaerobic digestion in electrochemical systems should consider the use of similar biogas compositions in the headspace to reflect real digestor conditions.

Our study provided a DIET co-culture with intermittently poised cathodes, as current interruption has been shown to be detrimental to a methanogenic bioelectrochemical reactor in which the cathode poised at −500mV (vs. SHE) was the sole source of electrons (Bretschger et al., 2015). Bretsgeer et al., observed a long recovery period (approx. 4 months) was required to reach similar methanogenesis levels as before the current interruption (Bretschger et al., 2015). In our system, the methanogen had access to more than one source of electrons at any time and yet they appear to favor cathodic H_2_, which leads to the demise of their syntrophic partner.

## Conclusion

This study investigated the effects of electrochemical stimulation on a DIET co-culture of *G. metallireducens* and *M. barkeri*. We found, for the first time, that under a biogas atmosphere, *M. barkeri* switched from DIET to methanogenesis with H_2_ from a cathode poised at −700mV as electron donor, independent of intermittent or continuous current additions. This new electron donor for the methanogen in the DIET consortia, destabilized the association, leading to significant decline in the *Geobacter* population. However, *Geobacter* appears robust and survives starvation periods resuming ethanol oxidation without significant lag. Overall low potential current addition was unfavorable to both DIET partners as apparent from final cell numbers as well as overall methane yield when compared to DIET co-cultures grown in similar reactors. Further studies with cathodes poised at higher potentials (−400 mV vs. SHE) are warranted to devise better protocols for biogas upgrading with inoculums from anaerobic digestors which could harbor potential DIET communities.

## Acknowledgements

This work was a contribution to a grant from the Innovationsfonden Denmark awarded to LDMO at Aarhus University (grant no. 4106–00017) and an ascending investigator grant by the Novo Nordisk Foundation awarded to AER at the University of Southern Denmark (grant no. NNF21OC0067353). We would like to thank Lasse Ørum-Smidt for lab assistance.

## References

Agneessens, L. M., Ottosen, L. D. M., Voigt, N. V., Nielsen, J. L., Jonge, N. de, Fischer, C. H., et al. (2017). IN-Situ Biogas Upgrading with Pulse H2 Additions: the Relevance of Methanogen Adaption and Inorganic Carbon Level. Bioresour. Technol. doi:10.1016/j.biortech.2017.02.016.

Aryal, N., Kvist, T., Ammam, F., Pant, D., and Ottosen, L. D. M. (2018). An overview of microbial biogas enrichment. Bioresour. Technol. 264, 359–369. doi:10.1016/j.biortech.2018.06.013.

Batlle-Vilanova, P., Puig, S., Gonzalez-Olmos, R., Vilajeliu-Pons, A., Balaguer, M. D., and Colprim, J. (2015). Deciphering the electron transfer mechanisms for biogas upgrading to biomethane within a mixed culture biocathode. RSC Adv. 5, 52243–52251. doi:10.1039/C5RA09039C.

Bauer, F., Persson, T., Hulteberg, C., and Tamm, D. (2013). Biogas upgrading - technology overview, comparison and perspectives for the future. Biofuels, Bioprod. Biorefining 7, 499–511. doi:10.1002/bbb.1423.

Beese-Vasbender, P. F., Grote, J.-P., Garrelfs, J., Stratmann, M., and Mayrhofer, K. J. J. (2015). Selective microbial electrosynthesis of methane by a pure culture of a marine lithoautotrophic archaeon. Bioelectrochemistry 102, 50–55. doi:10.1016/j.bioelechem.2014.11.004.

Bretschger, O., Carpenter, K., Phan, T., Suzuki, S., Ishii, S., Grossi-Soyster, E., et al. (2015). Functional and taxonomic dynamics of an electricity-consuming methane-producing microbial community. Bioresour. Technol. 195, 254–64. doi:10.1016/j.biortech.2015.06.129.

Chen, X., Ottosen, L. D. M., and Kofoed, M. V. W. (2019). How low can you go: Methane production of methanobacterium congolense at low CO 2 concentrations. Front. Bioeng. Biotechnol. 7, 34. doi:10.3389/fbioe.2019.00034.

Cheng, S., Xing, D., Call, D. F., and Logan, B. E. (2009). Direct biological conversion of electrical current into methane by electromethanogenesis. Environ. Sci. Technol. 43, 3953–3958. doi:10.1021/es803531g.

Da Costa Gomez, C. (2013). Biogas as an energy option: an overview. biogas Handb. Sci. Prod. Appl., 104–128. doi:10.1533/9780857097415.frontmatter.

De Vrieze, J., Hennebel, T., Boon, N., and Verstraete, W. (2012). Methanosarcina: the rediscovered methanogen for heavy duty biomethanation. Bioresour. Technol. 112, 1–9. doi:10.1016/j.biortech.2012.02.079.

Deutzmann, J. S., Sahin, M., and Spormann, A. M. (2015). Extracellular Enzymes Facilitate Electron Uptake in Biocorrosion and bioelectrosynthesis. MBio 6, 1–8. doi:10.1128/mBio.00496-15.Editor.

Ferguson, T. J., Mah, R. A., and Kuhn, D. A. (1983). Effect of H2-CO2 on Methanogenesis from Acetate or Methanol in Methanosarcina spp. Available at: https://www.ncbi.nlm.nih.gov/pmc/articles/PMC239386/pdf/aem00165-0060.pdf [Accessed September 14, 2019].

Fu, Q., Fukushima, N., Maeda, H., Sato, K., and Kobayashi, H. (2015). Bioelectrochemical analysis of a hyperthermophilic microbial fuel cell generating electricity at temperatures above 80 °C. Biosci. Biotechnol. Biochem. 79, 1200–1206. doi:10.1080/09168451.2015.1015952.

Garcia-Robledo, E., Ottosen, L. D. M., Voigt, N. V., Kofoed, M. W., and Revsbech, N. P. (2016). Micro-scale H2–CO2 Dynamics in a Hydrogenotrophic Methanogenic Membrane Reactor. Front. Microbiol. 7, 1276. doi:10.3389/fmicb.2016.01276.

Geppert, F., Liu, D., Van Eerten-Jansen, M. C. A. A., Weidner, E., Buisman, C. J. N., and ter Heijne, A. (2016). Bioelectrochemical Power-to-Gas: State of the Art and Future Perspectives. Trends Biotechnol. 34, 879–894. doi:10.1016/j.tibtech.2016.08.010.

Gregory, K. B., Bond, D. R., and Lovley, D. R. (2004). Graphite electrodes as electron donors for anaerobic respiration. Environ. Microbiol. 6, 596–604. doi:10.1111/j.1462-2920.2004.00593.x.

Hutten, T. J., Bongaerts, H. C. M., van der Drift, C., and Vogels, G. D. (1980). Acetate, methanol and carbon dioxide as substrates for growth of Methanosarcina barkeri. Antonie Van Leeuwenhoek 46, 601–610. doi:10.1007/BF00394016.

Kral, T. A., Brink, K. M., Miller, S. L., and McKay, C. P. (1998). Hydrogen consumptions by methanogens on the early earth. Orig. Life Evol. Biosph. 28, 311–319. doi:Doi 10.1023/A:1006552412928.

Lee, J. Y., Park, J. H., and Park, H. D. (2017). Effects of an applied voltage on direct interspecies electron transfer via conductive materials for methane production. Waste Manag. 68, 165–172. doi:10.1016/j.wasman.2017.07.025.

Lin, C., Wu, P., Liu, Y., Wong, J. W. C., Yong, X., Wu, X., et al. (2019). Enhanced biogas production and biodegradation of phenanthrene in wastewater sludge treated anaerobic digestion reactors fitted with a bioelectrode system. Chem. Eng. J. 365, 1–9. doi:10.1016/j.cej.2019.02.027.

Lovley, D. R. (1985). Minimum threshold for hydrogen metabolism in methanogenic bacteria. Appl. Environ. Microbiol. 49, 1530–1.

Lovley, D. R., Ueki, T., Zhang, T., Malvankar, N. S., Shrestha, P. M., Flanagan, K. A., et al. (2011). “Geobacter. The Microbe Electric’s Physiology, Ecology, and Practical Applications,” in Advances in Microbial Physiology (Academic Press), 1–100. doi:10.1016/B978-0-12-387661-4.00004-5.

Rosenbaum, M. A., Aulenta, F., Villano, M., and Angenent, L. T. (2011). Cathodes as electron donors for microbial metabolism: Which extracellular electron transfer mechanisms are involved? Bioresour. Technol. 102, 324–333. doi:10.1016/j.biortech.2010.07.008.

Rotaru, A.-E., Shrestha, P. M., Liu, F., Markovaite, B., Chen, S., Nevin, K. P., et al. (2014a). Direct Interspecies Electron Transfer between Geobacter metallireducens and Methanosarcina barkeri. Appl. Environ. Microbiol. 80, 4599–4605. doi:10.1128/AEM.00895-14.

Rotaru, A.-E., Shrestha, P. M., Liu, F., Shrestha, M., Shrestha, D., Embree, M., et al. (2014b). A new model for electron flow during anaerobic digestion: direct interspecies electron transfer to Methanosaeta for the reduction of carbon dioxide to methane. Energy Environ. Sci. 7, 408–415. doi:10.1039/C3EE42189A.

Rotaru, A. E., Yee, M. O., & Musat, F. (2021). Microbes trading electricity in consortia of environmental and biotechnological significance. Current Opinion in Biotechnology, 67, 119–129.

Rowe, A. R., Xu, S., Gardel, E., Bose, A., Girguis, P., Amend, J. P., et al. (2019). Methane-Linked Mechanisms of Electron Uptake from Cathodes by Methanosarcina barkeri. MBio 10, e02448–18. doi: 10.1128/mbio.02448-18.

Sharma, M., Varanasi, J. L., Jain, P., Dureja, P., Lal, B., Dominguez-Benetton, X., et al. (2014). Influence of headspace composition on product diversity by sulphate reducing bacteria biocathode. Bioresour. Technol. 165, 365–371. doi:10.1016/j.biortech.2014.03.075.

Shrestha, P. M., Malvankar, N. S., Werner, J. J., Franks, A. E., Elena-Rotaru, A., Shrestha, M.,… & Lovley, D. R. (2014). Correlation between microbial community and granule conductivity in anaerobic bioreactors for brewery wastewater treatment. Bioresource technology, 174, 306–310.

Siegert, M., Li, X.-F., Yates, M. D., and Logan, B. E. (2014). The presence of hydrogenotrophic methanogens in the inoculum improves methane gas production in microbial electrolysis cells. Front. Microbiol. 5, 778. doi:10.3389/fmicb.2014.00778.

Siegert, M., Yates, M. D., Spormann, A. M., and Logan, B. E. (2015). Methanobacterium Dominates Biocathodic Archaeal Communities in Methanogenic Microbial Electrolysis Cells. ACS Sustain. Chem. Eng. 3, 1668–1676. doi:10.1021/acssuschemeng.5b00367.

Smith, M. R., and Mah, R. A. (1978). Growth and Methanogenesis by Methanosarcina Strain 227 on Acetate and Methanol Downloaded from. Available at: http://aem.asm.org/ [Accessed January 17, 2019].

Stams, A. J. M., Sousa, D. Z., Kleerebezem, R., and Plugge, C. M. (2012). Role of syntrophic microbial communities in high-rate methanogenic bioreactors. Water Sci. Technol. 66, 352–362. doi:10.2166/wst.2012.192.

Van Eerten-Jansen, M. C. A. A., Heijne, A. Ter, Buisman, C. J. N., and Hamelers, H. V. M. (2012). Microbial electrolysis cells for production of methane from CO2: long-term performance and perspectives. Int. J. Energy Res. 36, 809–819. doi:10.1002/er.1954.

van Eerten-Jansen, M. C. A. A., Jansen, N. C., Plugge, C. M., de Wilde, V., Buisman, C. J. N., and ter Heijne, A. (2015). Analysis of the mechanisms of bioelectrochemical methane production by mixed cultures. J. Chem. Technol. Biotechnol. 90, 963–970. doi:10.1002/jctb.4413.

Van Eerten-Jansen, M. C. A. A., Veldhoen, A. B., Plugge, C. M., Stams, A. J. M., Buisman, C. J. N., and Ter Heijne, A. (2013). Microbial community analysis of a methane-producing biocathode in a bioelectrochemical system. Archaea 2013.

van Haandel, A., De Vrieze, J., Verstraete, W., and dos Santos, V. S. (2014). Methanosaeta dominate acetoclastic methanogenesis during high-rate methane production in anaerobic reactors treating distillery wastewaters. J. Chem. Technol. Biotechnol. 89, 1751–1759. doi:10.1002/jctb.4255.

Villano, M., Aulenta, F., Ciucci, C., Ferri, T., Giuliano, A., and Majone, M. (2010). Bioelectrochemical reduction of CO2 to CH4 via direct and indirect extracellular electron transfer by a hydrogenophilic methanogenic culture. Bioresour. Technol. 101, 3085–3090. doi:10.1016/j.biortech.2009.12.077.

Yee, M. O., Snoeyenbos-West, O. L. O., Thamdrup, B., Ottosen, L. D. M., and Rotaru, A.-E. (2019). Extracellular electron uptake by two Methanosarcina species. Front. Energy Res. 7. doi: 10.1101/458091.

Yee, M. O., & Rotaru, A. E. (2020). Extracellular electron uptake in Methanosarcinales is independent of multiheme c-type cytochromes. Scientific reports, 10(1), 1–12.

Zhao, Z., Zhang, Y., Quan, X., and Zhao, H. (2015a). Evaluation on direct interspecies electron transfer in anaerobic sludge digestion of microbial electrolysis cell. Bioresour. Technol. 200, 235–244. doi:10.1016/j.biortech.2015.10.021.

Zhao, Z., Zhang, Y., Wang, L., and Quan, X. (2015b). Potential for direct interspecies electron transfer in an electric-anaerobic system to increase methane production from sludge digestion. Sci. Rep. 5, 11094. doi:10.1038/srep11094.

Zhen, G., Kobayashi, T., Lu, X., and Xu, K. (2015a). Understanding methane bioelectrosynthesis from carbon dioxide in a two-chamber microbial electrolysis cells (MECs) containing a carbon biocathode. Bioresour. Technol. 186, 141–8. doi:10.1016/j.biortech.2015.03.064.

Zhen, G., Lu, X., Kobayashi, T., Kumar, G., and Xu, K. (2015b). Promoted electromethanosynthesis in a two-chamber microbial electrolysis cells (MECs) containing a hybrid biocathode covered with graphite felt (GF). Chem. Eng. J. 284, 1146–1155. doi:10.1016/j.cej.2015.09.071.

