## Supplementary materials for "The Fate of DIET Consortium exposed to continuous and intermittent electrochemical stimulation"

*Yee MO, Ottesen LDM, Rotaru AE, 2022*

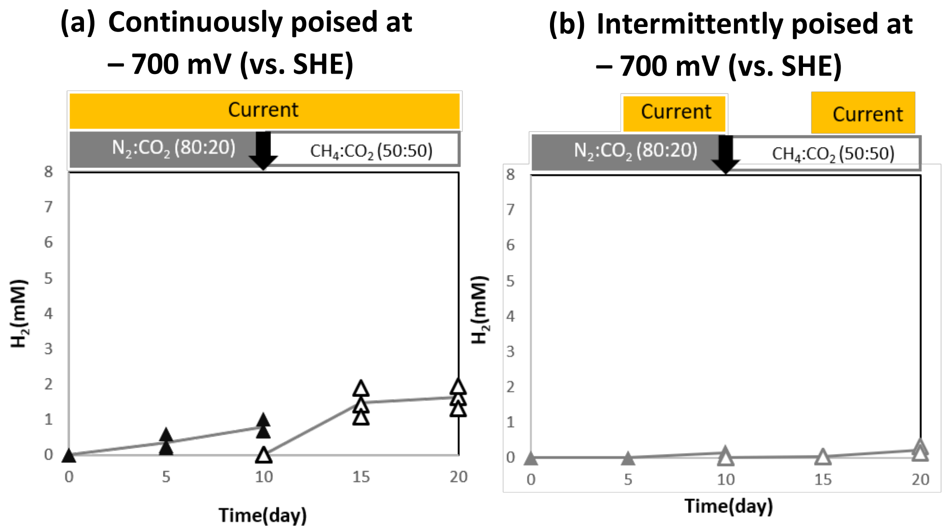


**Figure 4.** Hydrogen production in the abiotic bioelectrochemical reactors with a cathode poised at – 700 mV (vs. SHE) with (a) continuously poised cathodes (n=3) and (b) intermittently poised cathodes (n=2). The treatment strategy is depicted above the graph. The black arrow depicts time of gas exchange. The grey line is the average of replicates, closed symbols are measured under a headspace of $N_2$: $CO_2$ (80:20) and the open symbols are measured after flushing with $CH_4$: $CO_2$ (50:50)

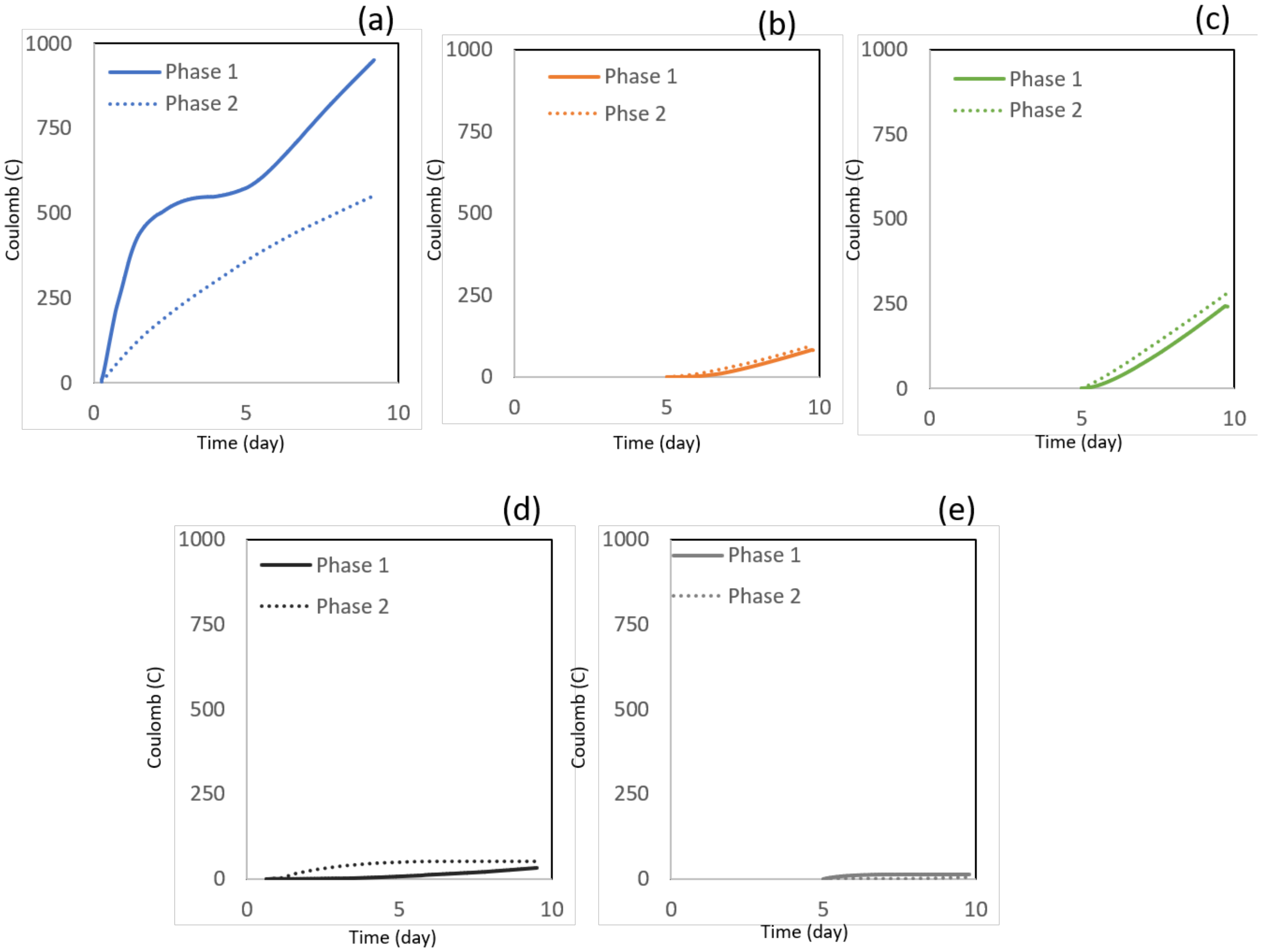


**Figure 2S.** Representative graphs of cumulative coulombs obtained over time in the bioelectrochemical reactors containing the co-culture of *G. metallireducens* and *M. barkeri* with a cathode poised at – 700 mV (vs. SHE) under different feed and current addition conditions, (a) Continuous feed and continuous current, (b) Continuous feed and intermittent current, (c) Intermittent feed or current. (d) Abiotic reactors with continuous electricity additions (e) Abiotic reactors with intermittent electricity additions
